# VCPA: genomic variant calling pipeline and data management tool for Alzheimer’s Disease Sequencing Project

**DOI:** 10.1101/327395

**Authors:** Yuk Yee Leung, Otto Valladares, Yi-Fan Chou, Han-Jen Lin, Amanda B Kuzma, Laura Cantwell, Liming Qu, Prabhakaran Gangadharan, Alzheimer’s Disease Sequencing Project (ADSP), William J Salerno, Gerard D. Schellenberg, Li-San Wang

**Affiliations:** Penn Neurodegeneration Genomics Center, Perelman School of Medicine at the University of Pennsylvania Richards Building, D101 3700 Hamilton Walk, Philadelphia, PA, USA.; Department of Pathology and Laboratory Medicine, Perelman School of Medicine at the University of Pennsylvania, PA, USA.; Human Genome Sequencing Center, Baylor College of Medicine, Houston, TX, USA.

## Abstract

**Summary:** We report VCPA, our SNP/Indel Variant Calling Pipeline and data management tool used for analysis of whole genome and exome sequencing (WGS/WES) for the Alzheimer’s Disease Sequencing Project. VCPA consists of two independent but linkable components: pipeline and tracking database. The pipeline is coded in Workflow Description Language and is fully optimized for the Amazon elastic compute cloud environment. This includes steps for processing raw sequence reads including read alignment, and all the way up to variant calling using GATK. The tracking database allows users to dynamically view the statuses of jobs running and the quality metrics reported by the pipeline. Users can thus monitor the production process and diagnose if any problem arises during the procedure. All quality metrics (>100 collected per processed genome) are stored in the database, thus facilitating users to compare, share and visualize the results. To summarize, VCPA is functional equivalent to the CCDG/TOPMed pipeline. Together with the dockerized database (also available as Amazon Machine Image), users can easily process any WGS/WES data on Amazon cloud with minimal installation.

**Availability:** VCPA is released under the MIT license and is available for academic and nonprofit use for free. The pipeline source code and step-by-step instructions are available from the National Institute on Aging Genetics of Alzheimer’s Disease Data Storage Site (http://www.niagads.org/VCPA).

**Supplementary information:** Supplementary data are available at *Bioinformatics* online.

## 1 Introduction

The Alzheimer’s Disease Sequencing Project (ADSP) is an integral component of the National Alzheimer’s Project Act (NAPA) towards a cure of Alzheimer’s Disease. ADSP will eventually analyze whole-genome sequencing (WGS) and whole-exome sequencing (WES) data from more than 20,000 late-onset Alzheimer’s Disease (AD) patients and cognitively normal elderly to finding new genetic variants associated with disease risk.

To ensure all sequencing data are processed following the best practices with consistency and efficiency, a common workflow was developed by the Genome Center for Alzheimer’s Disease (GCAD) in collaboration with ADSP. The workflow “Variant Calling Pipeline and data management tool” (VCPA), is used to process all the ADSP sequencing data. VCPA 1) is optimized for large-scale production of WGS and WES data, 2) includes a tracking database with web frontend for the user to track production process and review quality metrics 3) is implemented using the Workflow Description Language (WDL) for easier deployment and maintenance, 4) designed for the latest human reference genome build (GRCh38/hg38, version GRCh38DH) and follows best practices for WGS analysis with input from TOPMed (Trans-Omics for Precision Medicine) and CCDG (Centers for Common Disease Genomics), two other large sequencing programs supported by National Institutes of Health (NIH).

VCPA is composed of two independent but interoperable components: a tracking database (Figure 1A) with a web frontend (Figure 1B) and a SNP/indel calling pipeline (Figure 1C). The pipeline (available as an Amazon Machine Images (AMI): ami-82aa60f8) was optimized for automatic processing WGS/WES data in various file formats, from mapping sequence reads to the latest human reference genome (GRCh38/hg38) and variant calling. The tracking database (available as AMI: ami-acc840d3) was designed for monitoring the job status and recording quality metrics for each processed sample (Figure 1B). With a dynamic web interface of the database, researchers can easily compare, share and visualize all these individual level quality metrics.

**Figure 1:**
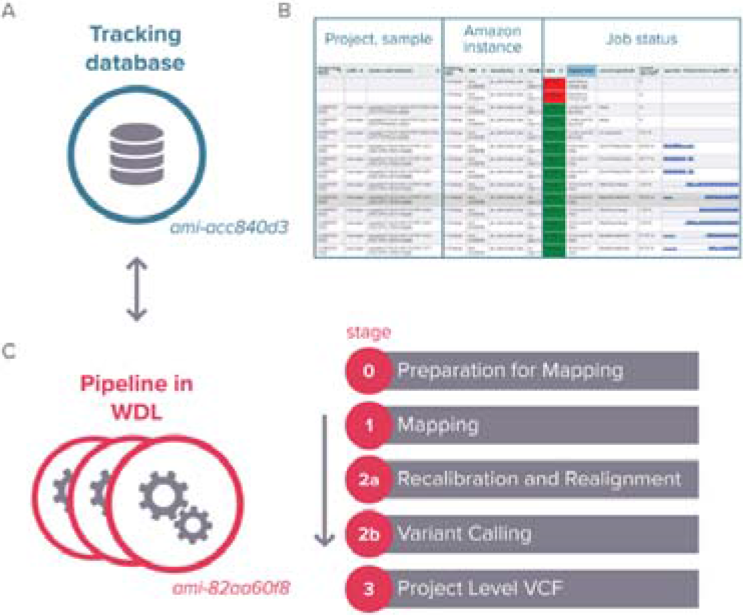
A) VCPA tracking database; B) Dynamic view of job status; C) VCPA Pipeline overview.

## 2 SNP/INDEL CALLING PIPELINE

The variant calling pipeline for the WGS (stages 1 and 2a) was developed in coordination with CCDG/TOPMed functional equivalent pipelines [2] and follows best practices of Germline Single Nucleotide Polymorphisms (SNPs) & Insertion/deletion (indel) Discovery for Genomic Analysis Toolkit (GATK) v3.7 [3]. VCPA pipeline is unique, as it accepts WGS or WES pair-end reads in FASTQ, BAM (binary sequence alignment map format), or CRAM (compressed BAM) formats with flow cell information and genomic regions for exome sequencing enrichment/capture kits. The processing workflow is modularized and consists of four stages (Figure 1C). If needed, the workflow can be configured to skip specific stages to reduce time and cost.

1. Stage 0 includes preparation steps for read mapping. For samples already mapped previously, PICARD [8] is used to roll back BAM files to uBAM (unaligned BAM) files.
2. Stage 1 generates BAM files. First, reads are mapped to GRCh38/hg38 using BWA-MEM [9], and duplicate reads are marked by BamUtil [10]. Next, BAM files are processed by Samblaster (adding MC and MQ tags to pair-end reads) [11] and sorted by genomic coordinates using SAMtools [12]. Finally, coverage statistics are computed using Sambamba [13].
3. Stage 2A performs local realignment near known indel sites (1000 Genome indels) and recalibration of base call quality scores using GATK v3.7 [14].
4. Stage 2B implements the GATK best practices steps for variant calling and annotation on SNPs and indels and generates genotype call files in genomic Variant Call Format (gVCF) for each sample individually. Quality metrics of called variants are computed using GATK [15].
5. Stage 3 combines gVCF files from multiple samples and performs joint genotype calling using GATK best practices. A project-level VCF file is generated.

For each project, the user starts by preparing a manifest file of sample IDs and sequencing read file locations. The manifest file is then uploaded into the tracking database at a head node server (the database can track multiple projects), and the user can submit processing jobs for the samples individually or by batches by command line from the head node. Job dependency and error checking are implemented in the workflow. Whenever multi-threading is supported by the third-party programs, jobs are run in parallel to expedite the process. A complete run of the pipeline produces ~200 files including analysis-ready quality scored binned read alignment BAM/CRAM files, annotated gVCF files, and more than 140 log files with data quality metrics and run information.

## 3 TRACKING DATABASE

The tracking database enables the user to monitor production status (Figure 1B) and review sequencing quality such as mapping percentage, depth coverage, and quality of called variants. The quality metrics pages are defined by projects and pipeline stages. All 113 quality metrics are collected during the pipeline execution and imported into the database, and are viewable through an interactive web frontend display.

The tracking database is built on a LAMP (Linux, Apache Httpd, MySQL and PHP) application stack using the SLIM-PHP framework. The application has a small memory and storage footprint, provides a RESTful API interface to the MySQL back-end, and supports password protection to restrict access. The tracking database is dockerized and can be installed on-site (off the cloud) if preferred.

## 4 USING VCPA ON AMAZON EC2

We evaluated our pipeline using the NA12878 sample from the Genome In A Bottle project using the hg38 high confident set [15]. The gVCF from running VCPA on GIAB sample were compared against the GIAB truth variant calls using hap.py [16]. Sensitivity/precision of VCPA calls were 0.999/0.994 for SNPs and 0.985/0.987 for indels respectively and comparable to TOPMed/CCDG workflows [2]. Using Amazon EC2 instance type r3.8xlarge (244.0 GiB, 32 vCPUs), we benchmarked running time on WGS data of 9 ADSP Discovery and Discovery Extension phase samples (average 78 million reads) with different paired-read length and file types: 1) three BAMs of 100 bp reads; 2) three BAMs of 150bp reads and 3) three CRAMs of 150bp reads. Average processing time per genome was 26.43, 22.68 and 21.20 core-hours respectively for the three configurations/file-types.

### Article short title

To conclude, VSPA is an efficient pipeline for processing high quality WGS/WES data on Amazon EC2 environment. VCPA is used for ADSP production and can track and receive information from >1,000 genome analysis runs simultaneously. Future plans include incorporating other variant calling pipelines such as xAtlas [17] and GATK4.

## Acknowledgements

We thank members of the Schellenberg and Wang labs, members from the GCAD and ADSP for their constructive input.

*Conflict of Interest:* none declared.

